# Predicting cognition using estimated structural and functional connectivity networks and artificial intelligence in multiple sclerosis

**DOI:** 10.1101/2025.03.07.642094

**Authors:** Ceren Tozlu, Dylan Ong, Christopher Piccirillo, Hannah Schwartz, Abhishek Jaywant, Thanh Nguyen, Keith Jamison, Susan A. Gauthier, Amy Kuceyeski

## Abstract

**Background:** Our prior work demonstrated that estimated structural and functional connectomes (eSC and eFC) generated using multiple sclerosis (MS) lesion masks and artificial intelligence (AI) models can predict disability as effectively as SC and FC derived from diffusion and functional MRI in MS. The goal of this study was to assess the ability of eSC and eFC in predicting baseline and 4-year follow-up cognition in MS patients.

**Methods:** One hundred seventy-one MS patients (age: 42.67±10.41, 74% females) were included. The Symbol Digit Modalities Test (SDMT), California Verbal Learning Test (CVLT), and Brief Visuospatial Memory Test (BVMT) were used to assess cognition. The Network Modification tool was performed to estimate SC, which was then used as an input to Krakencoder, an encoder-decoder model, to estimate FC. Ridge regression was performed to predict cognition using regional eSC and eFC, along with demographics and clinical information as well as conventional MRI metrics. Baseline cognition was added to the models that were used to predict the follow-up cognition. Spearman’s correlation (r) was used to assess the prediction accuracy.

**Results:** The highest accuracy was obtained when predicting follow-up SDMT using regional eSC or eFC (median r=0.58 for eSC and r=0.56 for eFC). Decreased eSC and eFC in the cerebellum and increased eFC in the default mode network were associated with lower follow-up SDMT scores. Baseline SDMT, clinical subtype, and age were the most important non-connectome metrics in predicting follow-up SDMT.

**Conclusions:** Our findings demonstrate that eSC and eFC derived from clinically acquired MRI and AI models can effectively predict cognition. The use of lesion-based estimates of connectome disruptions may potentially improve cognition-related individualized treatment planning.

## INTRODUCTION

Multiple Sclerosis (MS) is a chronic autoimmune disorder that causes ambulatory and cognitive impairment. Approximately 40-65% of MS patients suffer from varying levels of cognitive impairment^1^ including difficulties in attention^2^, processing speed^3^, and memory^4^. Neuroimaging biomarkers offer a valuable tool to identify patients who are more likely to develop cognitive impairment, enabling more personalized treatment plans to prevent or delay cognitive impairment.

Greater atrophy in the white matter (WM) and gray matter (GM)^5^ as well as higher lesion count and volume^6–8^ were shown to be associated with more cognitive impairment in MS. Recent studies have also shown that advanced MRI features derived from diffusion and functional MRI can be used to better understand neural circuitry underlying the cognitive impairment, as these features can provide information on how MS impacts brain networks, e.g. the brain’s structural connectomes (SC) and functional connectomes (FC)^9–12^. However, diffusion and functional MRI may not be accessible to most MS patients, as they are expensive and time-consuming for patients, and their processing requires researchers with a high level of domain expertise. Alternatively, our previous study has shown that the estimated SC and FC (eSC and eFC) obtained using only MS lesion masks derived from the clinical MRI perform similarly to observed SC and FC extracted from diffusion and functional MRI in predicting disease severity in people with MS^13^. However, the ability of eSC and eFC in predicting cognition has not yet been studied.

The goal of this study was to predict cognition at baseline and 4-year follow-up using eSC and eFC networks in a cohort of patients with MS. We first estimated SC networks via the Network Modification (NeMo) tool^14^ using the clinical MRI-derived lesion masks. Second, the Krakencoder^15^, a unified connectome tool that maps between structure and functional connectivity networks, was used to estimate FC networks from the eSC networks. Third, baseline and follow-up cognition were predicted using regional (i.e. node strength) eSC and eFC, which are calculated by taking the sum of the columns in the eSC or eFC matrix, respectively. We used regional connectomes rather than pairwise connectomes as our previous works showed that regional connectomes performed similarly or overperformed the pairwise connectomes in predicting disability as well as in classifying MS patients vs healthy controls^13,16^. We hypothesized that 1) eSC and eFC provide moderate accuracy in predicting cognition, 2) decreased eSC will be associated with lower cognitive scores, 3) both increased and decreased eFC will be associated with lower cognition as increased eFC might indicate an upregulation mechanism that delays cognitive impairment. Overall, our work provides evidence for the utility of clinical MRI-derived brain connectome estimates in 1) predicting cognition and 2) revealing neural circuitry underlying cognitive impairment in people with MS.

## MATERIALS AND METHODS

### Subjects

One hundred seventy-one patients with MS (age:42.67 ± 10.41, 74% females) with a diagnosis of clinically isolated syndrome (CIS) or MS (8 CIS, 158 relapsing remitting, 5 secondary progressive MS) were enrolled in a longitudinal imaging and cognitive study. Cognitive function was evaluated annually for up to four years utilizing The Brief International Cognitive Assessment for Multiple Sclerosis (BICAMS), which is composed of three assessments: Symbol Digit Modalities Test (SDMT), California Verbal Learning Test-II (CVLT-II) Immediate Recall (Total of Trials 1–5), and Brief Visuospatial Memory Test-Revised (BVMT-R) Immediate Recall (Total of Trials 1–3). The BICAMS is largely used in the MS field as it has been shown to be reliable and sensitive to cognitive changes and correlated with neuroimaging biomarkers such as gray matter volume in MS^17–19^. Among the BICAMS tests, SDMT is the most commonly used cognitive test in clinical care and research in the MS field.^20,21^ The cognitive tests were administered by research assistants trained by a neuropsychologist (A. J.) with regular meetings with a licensed clinical neuropsychologist (A. J.) and board-certified neurologist (S. G.) to ensure fidelity and accuracy of the application and results. The participants with a complete cognitive score were different for each cognitive test type at baseline and follow-up: n=167 for baseline SDMT, n=111 for follow-up SDMT, n=171 for baseline CVLT, n=114 for follow-up CVLT, n=171 for baseline BVMT, n=113 for follow-up BVMT (See Table 1). The 30% decrease in the number of subjects at the 4-year follow-up is probably due to the COVID-19 pandemic that happened in the second year of the study; subsequent year participation was impacted as well. Cognitive and neurological assessments were completed within an average of 14 ± 21 days of their annual MRI scan. Extended Disability Status Score (EDSS) was used to quantify disability and added to the prediction models. The inclusion criteria were (1) 2010 McDonald criteria^22^, (2) age ≥ 18 years and (3) already participants within our research repository [willing (and reconsented) to have an annual cognitive evaluation at the time of their annual MRI for a total of 4 years]. The participants were excluded if 1) they had contraindications to MRI, 2) the standard of care MRI scan was not completed within 45 days of the cognitive assessment, and 3) the patients developed other psychiatric disorders such as Alzheimer’s disease. Demographic data was collected on all patients (age, sex, clinical phenotype, and disease duration). The patients who were treated with Aubagio, Avonex, Betaseron, Copaxone, Tecfidera, IVIG, and Rebif were classified in the low efficacy treatment group, while the patients who were treated with Gilenya, Lemtrada, Ocrevus, Rituximab, and Tysabri were classified in the high efficacy treatment group. This study was approved by an ethical standards committee on human experimentation, and written informed consent was obtained from all patients.

**Table 1:** Patient characteristics. Demographics and clinical information for people with MS. The median [1st, 3rd quartile] was used for the continuous metrics, and the count was used for the categorical variables such as sex, and disease type. The demographic and imaging metrics at baseline were given for 171 patients. The number of patients with a complete SDMT, BVMT, and CVLT at baseline and follow-up was given in the last 3 rows of the table. The p-values comparing the baseline and follow-up cognitive scores were given in the last column. A paired Wilcoxon rank sum test was used to compare the baseline and follow-up cognitive scores.

| Variable | Baseline | 4 Year Follow-up | p-value |
| --- | --- | --- | --- |
| Age (years) | 41 [35,51] | - |  |
| Sex |  | - |  |
| Female | 126 |  |  |
| Male | 45 |  |  |
| Disease Type |  | - |  |
| CIS | 8 |  |  |
| RRMS | 158 |  |  |
| SPMS | 5 |  |  |
| Treatment Type |  | - |  |
| None | 1 |  |  |
| Low Efficacy | 77 |  |  |
| High Efficacy | 93 |  |  |
| Disease Duration (years) | 9.8 [5.15, 14.2] | - |  |
| EDSS | 1 [0, 2] | - |  |
| Cortical Thickness (mm) | 2.42 [ 2.37, 2.48] | - |  |
| Thalamic Volume (mm<sup>3</sup>) | 9.10 x 10<sup>-3</sup> [ 8.58 x 10<sup>-3</sup>, 9.58 x 10<sup>-3</sup>] | - |  |
| Lesion Volume (mm<sup>3</sup>) | 2200 [882.2, 6592.8] |  |  |
| SDMT | 58 [49, 64] (n=167) | 58 [52, 66] (n=111) | 0.38 |
| CVLT | 55 [46, 60] (n=171) | 59 [ 52, 66.75] (n=114) | <0.01 |
| BVMT | 25 [20.5, 29] (n=171) | 28 [23, 31] (n=113) | 0.003 |

### Image acquisition, processing, and connectome extraction

Imaging was performed on a 3T Magnetom Skyra scanner (Siemens Medical Solutions USA, Malvern, PA) using a product twenty-channel head/neck coil. The MRI protocol consisted of sagittal 3D T1-weighted (T1w) sequence for anatomical structure, 2D T2-weighted (T2w) fast spin echo, and 3D T2w fluid-attenuated inversion recovery (FLAIR) sequences for lesion detection, and gadolinium-enhanced 3D T1w sequence for acute lesion identification. A detailed description of the imaging protocol: (1) 3D sagittal T1w MPRAGE: Repetition Time (TR)/Echo Time (TE)/Inversion Time (TI)=2300/2.3/900 ms, flip angle (FA)=8°, GRAPPA parallel imaging factor (R)=2, voxel size=1.0×1.0×1.0 mm3; (2) 2D axial T2-weighted (T2w) turbo spin echo: TR/TE=5840/93 ms, FA=90, turbo factor=18, R=2, number of signal averages (NSA)=2, voxel size=0.5×0.5×3 mm3; (3) 3D sagittal fat-saturated T2w fluid attenuated inversion recovery (FLAIR) SPACE: TR/TE/TI=8500/391/2500 ms, FA=90°, turbo factor=278, R=4, voxel size=1.0×1.0×1.0 mm3; (4) axial 3D multi-echo GRE sequence for QSM: axial field of view (FOV)=24 cm, TR/TE1/δTE=48.0/6.3/4.1 ms, number of TEs=10, FA=15◦, R=2, voxel size=0.75×0.93×3 mm3, scan time=4.2 min.

### Lesion mask creation

The WM hyperintensity lesion masks were created by running the T2 FLAIR images through the Lesion Segmentation Tool (LST) and were further manually edited as needed. T2 FLAIR-based lesion masks were transformed to the individual’s T1 native space using the inverse of the T1 to GRE transform and trilinear interpolation. Individual T1 images were then normalized to MNI space using FSL’s linear (FLIRT) and non-linear (FNIRT) transformation tools (http://www.fmrib.ox.ac.uk/fsl/index.html); transformations with trilinear interpolation were then applied to transform the native anatomical space T2FLAIR lesion masks to MNI space. The transformations were concatenated (T2FLAIR to T1 to MNI) to minimize interpolation effects. Lesions were manually inspected after the transformation to MNI space to verify the accuracy of the coregistration.

### Estimation of the Structural and Functional Connectivity Networks

The NeMo Tool^14^ was applied to each subject’s MNI-space T2 FLAIR lesion mask to calculate eSC using a reference database of SCs from 420 unrelated HC (206 female, 214 male, 28.7 ± 3.7 years) from the from the Human Connectome Project Young Adult (HCP-YA) dataset (See Supplementary Document for more details). The NeMo Tool first identifies streamlines in the reference database (obtained using the deterministic [sd-stream] tractography with MRtrix3^23,24^) that pass through the lesion mask provided and removes them from further calculations. It then calculates the eSC by taking the sum of the SIFT2 weights of remaining streamlines connecting pairs of regions, divided by the sum of the two regions’ volumes averaged over the 420 individuals in the reference set. Regional eSC was calculated as the sum of the columns in the eSC matrix; the diagonal of the eSC matrix is zero.

After obtaining the eSC matrices with 3 different parcellations (FreeSurfer 86, Shen 268, and coco439) and 2 different tractography methods (probabilistic and deterministic) via the NeMo tool, the Krakencoder^15^ was applied to estimate FC from these eSC. The Krakencoder is an encoder-decoder model that was trained using healthy connectomes (HCP-YA) to bidirectionally map between SC and FC across different atlases and processing choices. Various eFC types including high vs band pass filtering and with vs without global signal regression were obtained via the Krakencoder. In this study, we used the eFC with high pass filtering and global signal regression similar to our most recent paper on MS^25^. Regional eFC was calculated by taking the sum of each of the columns in the eFC after removing the negative entries and assigning zero to the diagonal.

As required by the Krakencoder, the eSC matrices were obtained with 3 different parcellations (FreeSurfer 86, Shen 268, and coco439). However, the prediction of cognition was performed with the regional eSC and eFC constructed using FreeSurfer 86 atlas containing 68 cortical and 18 subcortical/cerebellar regions^26^. We used the same FreeSurfer atlas as our previous works that performed connectome-disability mapping in MS^13,16,25,27,28^ so we can compare the findings of this study with our previous results.

### Univariate analysis

Demographic and clinical variables were presented in Table 1 using the median [1st, 3rd quartile] for the continuous metrics, and using the count for the categorical variables such as sex, and disease type. A paired Wilcoxon rank-sum test was used to test whether there was a significant difference between baseline and follow-up cognition. Student’s t-test was used to identify the amplitude of the difference in pairwise eSC in MS patients vs SC in healthy young adults as well as pairwise eFC in MS patients vs FC in healthy young adults from the Human Connectome Project-Young Adult database. The statistics from this t-test were represented at pairwise and region levels in Figure 2. All statistical analyses were performed, and graphs were created using R version 3.4.4 and Matlab version R2020a.

### Regression analysis

Linear regression with ridge regularization was performed to predict cognition (SDMT, CVLT, and BVMT separately) using different combinations of demographics, clinical information, conventional MRI metrics (cortical thickness and normalized thalamic volume), regional eSC, and regional eFC. Baseline cognition scores were added to the model predicting follow-up cognition. All features used in each model are described below and represented in Figure 1:

1. Model I: Regional eSC, demographics, and clinical information as well as conventional MRI metrics (sex, age, race, disease duration, clinical phenotype, EDSS, cortical thickness, and normalized thalamic volume)
2. Model II: Regional eFC, demographics, and clinical information as well as conventional MRI metrics (sex, age, race, disease duration, clinical phenotype, EDSS, cortical thickness, and normalized thalamic volume)
3. Model III: Regional eSC and eFC, demographics and clinical information as well as conventional MRI metrics (sex, age, race, disease duration, clinical phenotype, EDSS, cortical thickness, and normalized thalamic volume)

**Figure 1:**
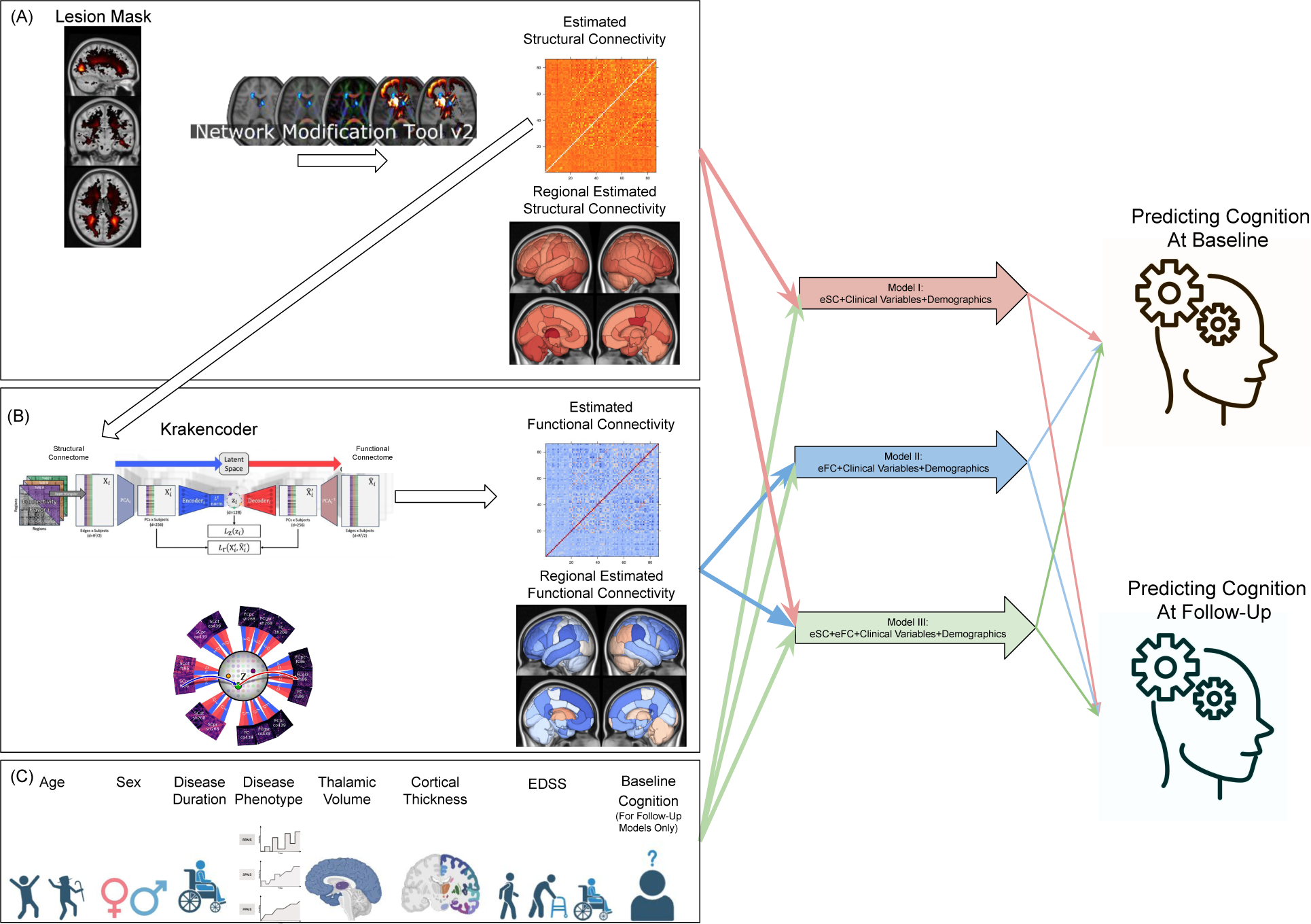
Workflow of the study. (A) eSC matrix was generated using the lesion masks derived from T2 FLAIR images and the Network Modification tool. (B) eSC matrix was used as an input to the Krakencoder to generate eFC matrices for each MS patient. The Krakencoder was trained using SC and FC matrices from young healthy adults (Human Connectome Project – Young Adult dataset). (C) Data including demographics and clinical information as well as conventional MRI metrics were added to the models based on eSC and eFC. Ridge regression was performed to predict cognition at baseline and follow-up. Spearman’s correlation coefficient between the original and predicted cognitive scores was used to evaluate the prediction accuracy of the models.

The models were trained with outer and inner loops of k-fold cross-validation (k = 5) to optimize the hyperparameters and test model performance. The inner loop (repeated over 5 different partitions of the training dataset only) optimized the set of hyperparameters that maximized validation set accuracy. A model was then fitted using the entire training dataset and those optimal hyperparameters, which were then assessed on the hold-out test set from the outer loop. The outer loop was repeated for 100 different random partitions of the data. The median of the distribution of Spearman’s correlations (overall 5 folds × 100 iterations = 500 test sets) was calculated as a summary of model performance. The prediction accuracy distributions for different models were compared using a permutation test^29^ (1000 permutations) where p-values were calculated as the number of permutations that had a difference in means bigger than the original difference. P-values were considered significant when p < 0.05 after BH correction for multiple comparisons ^30^ (Benjamini and Hochberg, 1995). The strength of the relationship of eSC and eFC with cognition was identified using the mean feature weights (the beta parameter coefficients) over all 500 models (100 partitions of the data into 5 folds). Regional variable importance metrics were also summarized at a functional network level by assigning each of the 68 cortical regions to one of seven canonical functional networks^31^; subcortex and cerebellum were also added as their own networks.

Additionally, Models I and II were also repeated using the pairwise eSC and eFC instead of regional eSC and eFC, respectively using 50 different random partitions of the data. The prediction accuracy of the pairwise models was presented in the Supplementary Document.

## RESULTS

### Patient Characteristics

The study participants were relatively young with a median age of 41, the majority having relapsing-remitting MS (158 out of 171), and approximately three-quarters being female, reflecting the typical demographics of the MS population (Table 1). The study participants had a minimal disability with a median EDSS of 1 and a small number of the participants had cognitive impairment (22%, 12%, and 26% of participants had cognitive impairment at baseline based on a norm-referenced z-score < −1 on the SDMT, CVLT, and BVMT, respectively). In participants who had both baseline and follow-up cognition scores, cognitive measurements either remained the same or slightly increased between the baseline year and year 4, which can be explained by the low percentage of participants with cognitive impairment at baseline and the fact that most patients (54%) were treated with high efficacy treatments.

### eSC and eFC in MS patients compared to health controls

The impact of the lesions on the structural and functional connectomes were shown in Figure 2. The statistics from the Student’s t-test was used to compare the eSC and eFC matrices in people with MS with the SC and FC in healthy young adults from the Human Connectome Project – Young Adults database. As expected, the people with MS showed decreased eSC due to MS lesions compared to healthy young adults. The most impacted structural networks were between subcortex and dorsal attention. MS patients showed decreased eFC between ventral attention and somatomotor cortex and increased eFC between the cerebellum and dorsal attention compared to healthy young adults.

**Figure 2:**
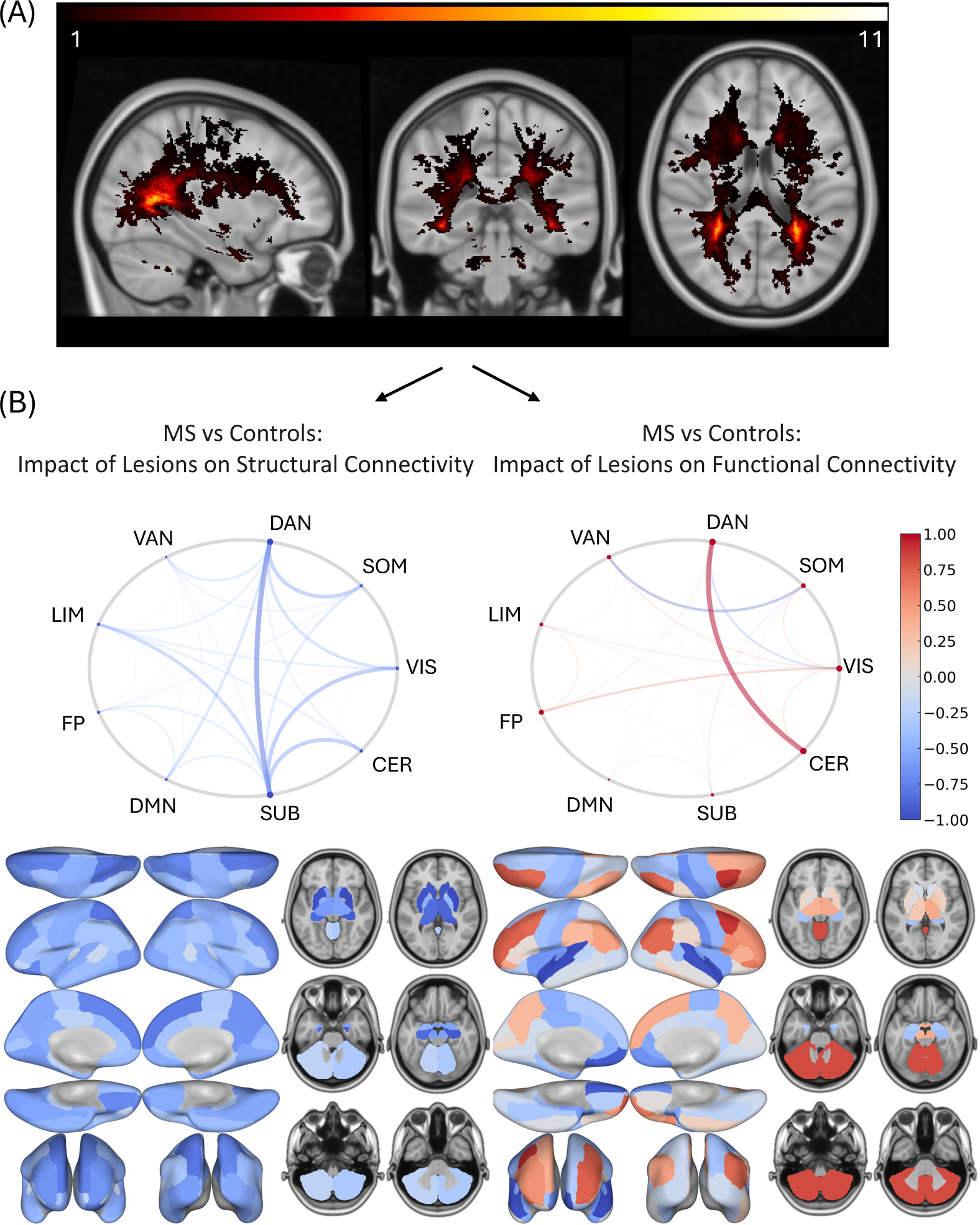
Impact of MS lesions on the structural and functional connectomes. (A) The average lesion masks in people with MS. Color indicates the number of individuals that had a lesion in that voxel. (B) The statistics from the Student’s t-test comparing the pairwise and regional eSC and eFC matrices in people with MS compared to healthy young adults from the Human Connectome Project – Young Adults database. For the circle plot, each of the 86 regions was assigned to one of nine networks, including the seven Yeo functional networks, a subcortical, and a cerebellar network. DAN = dorsal attention; VAN = ventral attention; LIM = limbic; FP = fronto-parietal; DMN = default-mode network; SUB = subcortex; CER = cerebellum; VIS = visual; SOM = somatomotor. Red colors represent increased connections in people with MS, while blue colors represent decreased connections in people with MS compared to healthy young adults.

### Model accuracy in predicting cognition

Figure 3 shows the accuracy of 3 different models in predicting baseline and 4-year follow-up cognitive scores. The Spearman’s correlation between the observed and predicted cognitive score was used to assess the models’ accuracy. The highest prediction accuracy was obtained when predicting 4-year follow-up SDMT. The eSC and eFC models performed equally well in predicting 4-year follow-up SDMT (median r=0.58 for eSC and r=0.56 for eFC). Finally, the accuracy of the models containing the pairwise eSC and eFC matrices performed worse than their regional counterparts (See Supplementary Figure 1).

**Figure 3:**
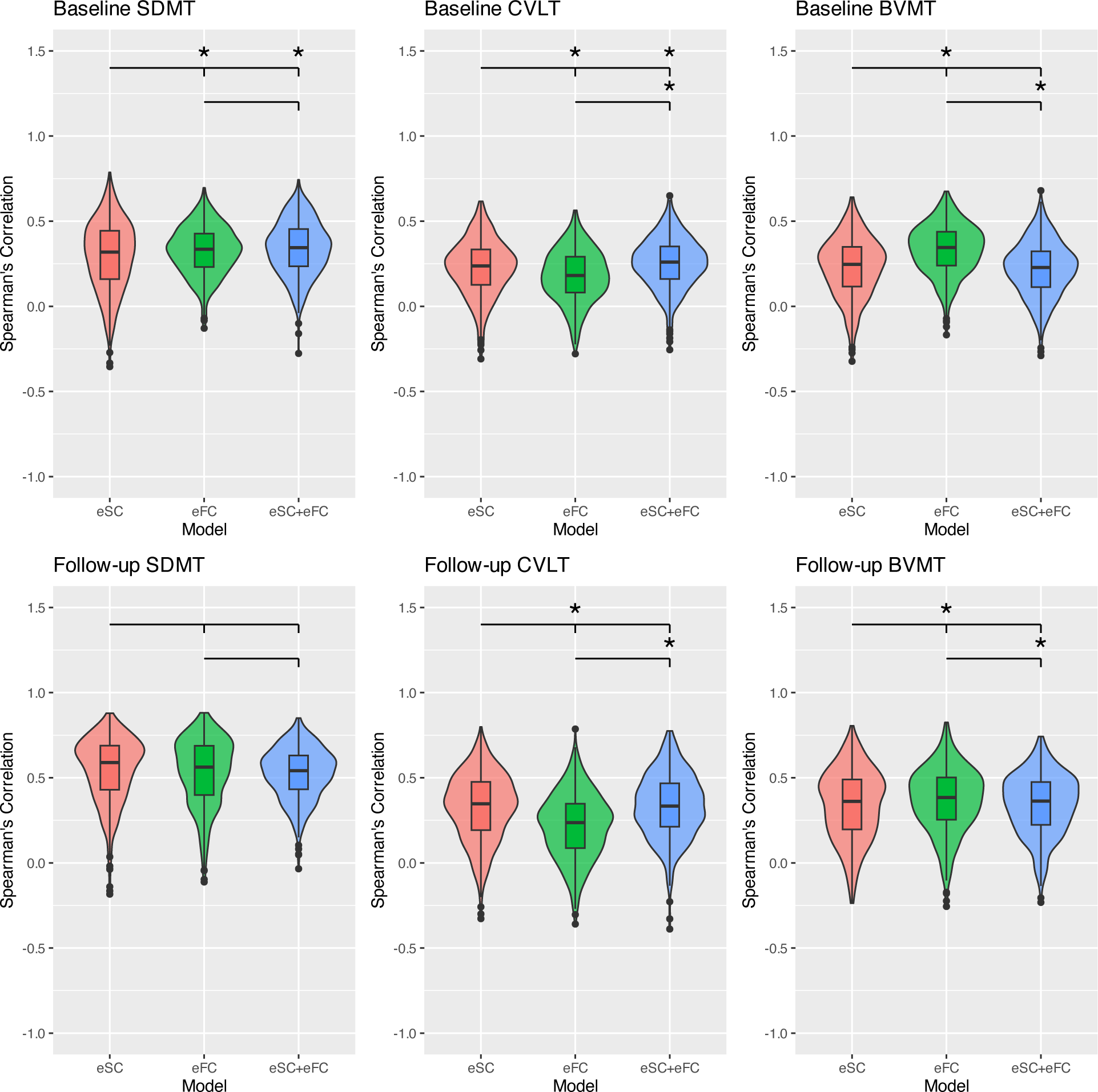
Accuracy in predicting baseline and follow-up cognition. Distributions of accuracy (Spearman’s correlation between the observed and predicted values) for models predicting cognitive metrics at baseline SDMT, BVMT, and CVLT (top row) and 4-year follow-up SDMT, BVMT, and CVLT (bottom row). A permutation test was used to compare each pair of models’ performances; * indicates permutation-based p < 0.05 after multiple comparison correction.

### Feature Importance

As the highest prediction performances were obtained when predicting 4-year follow-up SDMT, feature importance (i.e. the beta estimates from the ridge regression) from these models are presented in Figure 4. As expected, baseline SDMT score was the most important variable in predicting follow-up SDMT, followed by normalized thalamic volume. All models that included regional eSC and eFC metrics showed that the eSC and eFC in the cerebellum were the most important features in predicting follow-up SDMT, where decreased eSC and eFC in the cerebellum were associated with lower follow-up SDMT. Additionally, the model based only on eFC indicated decreased eFC in the DMN and to a slightly lesser extent frontoparietal, visual, and limbic networks and increased eFC in dorsal and ventral attention networks was associated with better follow-up SDMT.

**Figure 4:**
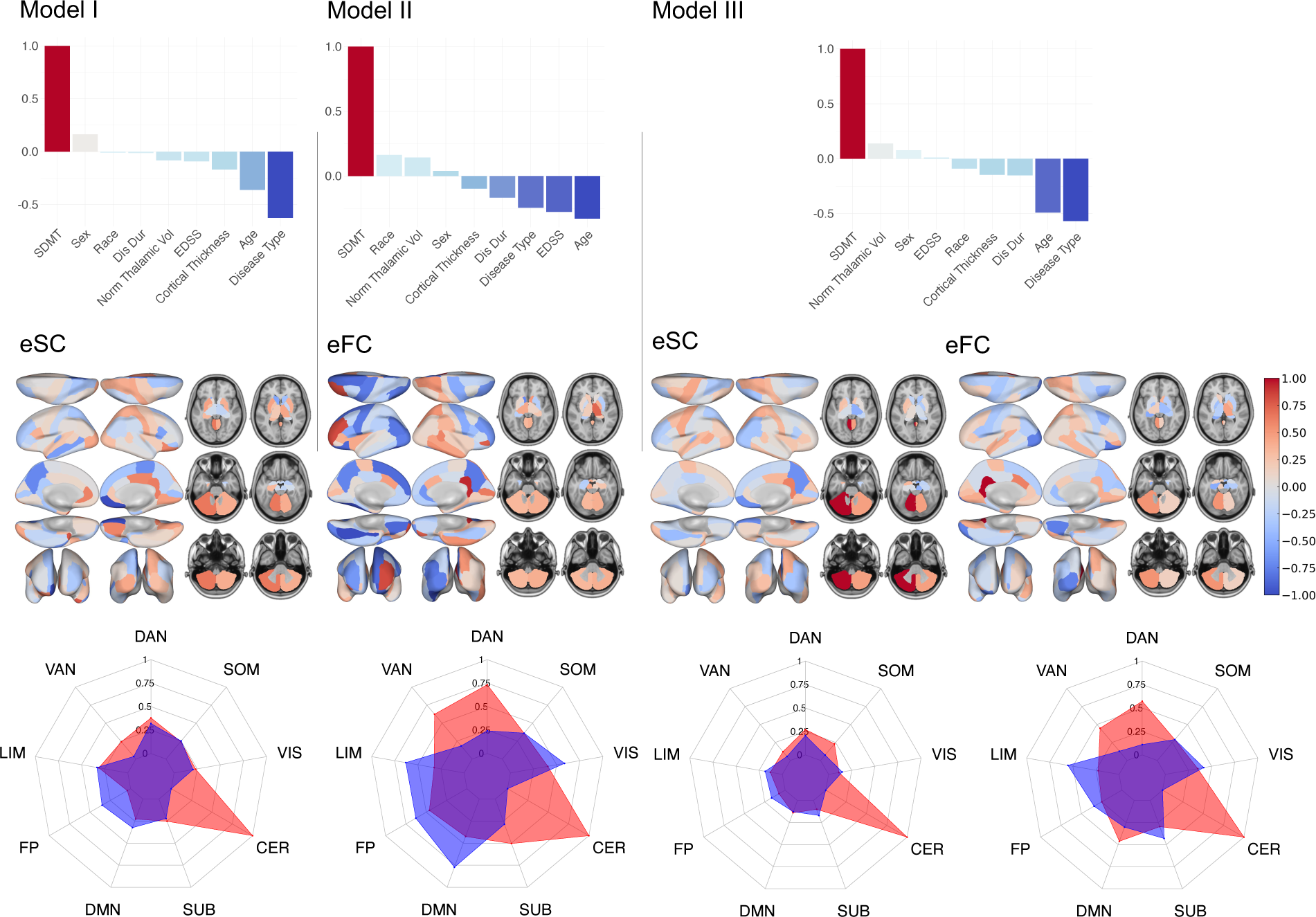
Variable importance in predicting the follow-up SDMT. Variable importance (normalized ridge regression beta estimates) of regional eSC, regional eFC, demographics, and clinical information as well as conventional MRI metrics calculated from 3 different models to predict year 4 follow-up SDMT. The importance of the demographics and clinical information as well as conventional MRI metrics are shown via bar plots. Positive importance values (red) can be interpreted as increased values at baseline are related to higher 4-year SDMT, while negative importance values (blue) can be interpreted as decreased values at baseline are related to higher 4-year SDMT. The importance of the regional eSC and eFC was presented using brain plots.

## DISCUSSION

Our study aimed to demonstrate the ability of estimated brain connectomes, combined with demographics and clinical information as well as conventional MRI metrics, to predict baseline and follow-up cognition in MS patients. Overall, our findings indicated that estimated structural and functional connectomes can moderately predict cognition in MS patients. Additionally, our results also showed that (1) all models achieved the highest accuracy when predicting the follow-up SDMT compared to other cognitive scores, (2) the model containing the eSC and eFC performed moderately and equally well in predicting the follow-up SDMT, and (3) increased eSC and eFC in the cerebellum, decreased eFC in the DMN and increased eFC in attention networks were predictive of higher 4-year follow-up SDMT.

### eSC and eFC can moderately predict cognition in MS

Our previous study has shown that eSC and eFC can predict the ambulatory disability level significantly better than SC and FC metrics derived from diffusion and functional MRI collected directly in MS patients^13^. The same study also showed that the prediction performance of both estimated and observed SC and FC in predicting disability was moderate. Expanding this work, we here provided evidence that eSC and eFC can also moderately predict cognition. The highest prediction accuracy in our study was above 0.5, which is a higher or similar performance compared to other studies that predicted cognition in MS patients^10^ and healthy individuals.^32–36^ Models demonstrated moderate performance in predicting both baseline and follow-up cognition. However, their performance was notably better in predicting follow-up cognition than baseline cognition. It is very likely due to the inclusion of baseline cognition performance as a predictor in the follow-up cognition models; there is generally a high correlation between baseline and follow-up cognition (Spearman’s correlation coefficients of 0.76 for SDMT [p-value < 2.2e-16], 0.55 for CVLT [p-value = 1.755e-13], and 0.70 for BVMT [p-value = 9.941e-16]).

### Cerebellar connections as an important biomarker of motor and cognitive impairment in MS

Structural and functional connections involving the cerebellum have been shown as important biomarkers of disability in MS^13,37–39^. Our previous research demonstrated that the disconnection resulting from paramagnetic rim lesions (PRLs) or non-PRLs in the cerebellum was among the strongest predictors of severe disability in MS^39^. Additionally, we also found that decreased eSC and increased eFC were linked to higher levels of ambulatory disability in MS, emphasizing the cerebellum’s critical role in disability in people with MS^13^. While the cerebellum’s role in motor function has long been established, recent works have highlighted its involvement in cognitive processes such as executive functioning, language, and spatial cognition.^40,41^ MS studies have further shown that cerebellar abnormalities were not only associated with motor impairment but also with cognitive decline.^42–46^ Increased cerebellar atrophy and cerebellar T1 lesion volume were the strongest predictors of lower SDMT^47^. Another study also reported increased functional activation in the cerebellum when performing SDMT tasks during an fMRI scan, underscoring the cerebellum’s pivotal role in information processing speed and supporting our results with univariate and multivariate approaches^48^.

### Maladaptation mechanism in the DMN

Our results also suggested that increased eFC in the DMN was associated with lower follow-up SDMT, potentially indicating a maladaptation mechanism within the DMN. Previous studies have reported conflicting results regarding the association of the FC in the DMN and cognition in MS. These discrepancies may stem from variations in methodological approaches, such as analyzing different types of connectivity (e.g., within-network, between-network, or connectivity with the rest of the brain) or investigating the DMN as a whole versus dividing it into specific regions of interest. While decreased FC in the anterior components of the DMN was shown in MS patients with a cognitive impairment^49,50^, increased FC of the DMN with the rest of the brain was observed in MS patients with cognitive impairment compared to the patients without cognitive impairment and healthy controls^51^. Our approach uses a similar approach with the latter study (e.g. FC of the DMN with the rest of the brain), which also supports the hypothesis of a maladaptation mechanism related to the DMN in MS patients.

### Advantages of using the estimated connectomes

One of the major advantages of using estimated connectomes rather than observed connectomes is accessibility. In a general clinical setting with monetary, time, and expertise constraints, taking a simple MRI scan to obtain lesion masks is much more practical than collecting and processing advanced diffusion and functional MRI scans. Additionally, patient conditions like edema and inflammation can add noise to dMRI and fMRI measurements; eSC and eFC generated by NeMo and Krakencoder will not suffer from these decreases in SNR^13^. Estimated connectomes that use lesion information only may also allow isolation of the effect of pathology on the connectomes, ignoring other population-level variability that is not relevant to patient performance. Finally, using the NeMo tool and Krakencoder to generate eSC and eFC from clinical MRI means large, already existing datasets can be utilized for connectome-behavior mapping. In conclusion, using eSC and eFC can be advantageous over their observed counterparts due to better ease of access, more consistent data, and larger quantities of training data.

### Limitations and future works

One major limitation of studies using eSC and eFC metrics is the reliance on healthy individuals’ SC (in the NeMo tool) or healthy individuals’ SC-FC relationships (in the Krakencoder), which may be different in a patient cohort that has brain lesions. However, a previous study has shown that the disconnectivity metrics are similar (a correlation of 0.866 ± 0.066) when the tractography was used from age-matched controls vs young adults in the Human Connectome Project^52^. Another limitation is that the potential gray matter lesions and other pathologies in the white matter were not considered by the NeMo tool as only the white matter lesion mask is considered in the calculation of the eSC. On the other hand, our previous study has shown that the eSC and eFC perform similarly to the SC and FC, providing evidence that other pathologies may not contribute as much to the disability as the white matter lesions themselves. Another limitation of our study was that most patients were relatively young (median age was 41), did not have cognitive impairment at baseline, and were in the early stage of the disease, with about 92.5% of the research subjects having RRMS. As a result, on average, participants’ performance on the cognitive tests either remained the same or slightly increased, which may be due to patients receiving highly effective disease-modifying therapies, which are associated with slower progression in cognition as well as ambulatory progression^53^. Therefore, a future study including a higher number of older and progressive patients is needed to better generalize the findings to all MS types. Another limitation was that the current study uses the cognition scores at a 4-year follow-up, which may be too brief a period to capture cognitive changes. Therefore, a future study with a greater time difference between the baseline and follow-up is needed to provide greater insights into the ability of the eSC and eFC to predict cognition, leading to more personalized care with long-term impact. Lastly, the number of participants decreased by 30% at a 4-year follow-up due to the COVID-19 pandemic, which began in the second year of the study and affected subsequent years. Participants who dropped out of the study during the pandemic may have been impacted more by the pandemic and experienced greater cognitive impairment.

### Conclusion

In summary, our work demonstrated the feasibility and utility of using lesion-informed connectome estimates in predicting various cognitive domains and understanding the neural circuitry of cognitive impairment in MS. Our study showed that lesion-informed connectome estimates extracted from clinically acquired, conventional MRI can moderately predict follow-up cognition. This work highlights the potential for using connectome mapping in the clinic to make personalized treatment plans aimed at preserving cognition in people with MS.

## Acknowledgments

C.T. designed the study, performed the statistical analysis, and wrote the article.

D.O. performed the statistical analysis and wrote the article.

C.P. performed the statistical analysis and wrote the article.

H.S. collected the data and edited the article.

A.J. supervised the study and edited the article.

K.J. generated the estimated FC metrics, helped with the co-registration of the lesion masks, and edited the article.

S.G. designed and supervised the study, collected the data, and edited the article.

A.K. designed and supervised the study and edited the article.

## Data/code availability statement

The Network Modification tool can be reached via: https://kuceyeski-wcm-web.s3.us-east-1.amazonaws.com/upload.html. The codes for Krakencoder can be found at https://github.com/kjamison/krakencoder. The ridge regression codes that were used in this study are publicly available here: https://github.com/cerent/BICAMS-NeMo-Krakencoder. The deidentified data (the estimated connectivity matrices) that support the findings of this study are available upon reasonable request from the corresponding author.

## Ethics statement

All studies were approved by an ethical standards committee of Weill Cornell Medicine on human experimentation, and written informed consent was obtained from all patients.

## References

1. Sumowski, J. F. et al. Cognition in multiple sclerosis: State of the field and priorities for the future. Neurology 90, 278–288 (2018).

2. Jansen, D. A. & Cimprich, B. Attentional impairment in persons with multiple sclerosis. J. Neurosci. Nurs. J. Am. Assoc. Neurosci. Nurses 26, 95–102 (1994).

3. van Geest, Q. et al. Information processing speed in multiple sclerosis: Relevance of default mode network dynamics. NeuroImage Clin. 19, 507–515 (2018).

4. Basso, M. R. et al. Memory in multiple sclerosis: A reappraisal using the item specific deficit approach. Neuropsychology 35, 207–219 (2021).

5. Rimkus, C. de M., et al. Atrophy Patterns in Patients With Multiple Sclerosis With Cognitive Impairment, Fatigue, and Mood Disorders. Neurology 103, e210080 (2024).

6. Engl, C. et al. Cognitive impairment in early MS: contribution of white matter lesions, deep grey matter atrophy, and cortical atrophy. J. Neurol. 267, 2307–2318 (2020).

7. Harrison, D. M. et al. Association of Cortical Lesion Burden on 7-T Magnetic Resonance Imaging With Cognition and Disability in Multiple Sclerosis. JAMA Neurol. 72, 1004–1012 (2015).

8. Rovaris, M. et al. Cortical/subcortical disease burden and cognitive impairment in patients with multiple sclerosis. AJNR Am. J. Neuroradiol. 21, 402–408 (2000).

9. Buyukturkoglu, K. et al. Classifying multiple sclerosis patients on the basis of SDMT performance using machine learning. Mult. Scler. J. (2020) doi:10.1177/1352458520958362.

10. Fuchs, T. A. et al. Functional Connectivity and Structural Disruption in the Default-Mode Network Predicts Cognitive Rehabilitation Outcomes in Multiple Sclerosis. J. Neuroimaging 30, 523–530 (2020).

11. Fuchs, T. A. et al. Preserved network functional connectivity underlies cognitive reserve in multiple sclerosis. Hum. Brain Mapp. hbm.24768 (2019) doi:10.1002/hbm.24768.

12. Kletenik, I. et al. Multiple sclerosis lesions that impair memory map to a connected memory circuit. J. Neurol. 270, 5211–5222 (2023).

13. Tozlu, C., Jamison, K., Gu, Z., Gauthier, S. A. & Kuceyeski, A. Estimated connectivity networks outperform observed connectivity networks when classifying people with multiple sclerosis into disability groups. NeuroImage Clin. 32, 102827 (2021).

14. Kuceyeski, A., Maruta, J., Relkin, N. & Raj, A. The Network Modification (NeMo) Tool: Elucidating the Edect of White Matter Integrity Changes on Cortical and Subcortical Structural Connectivity. Brain Connect. 3, 451–463 (2013).

15. Jamison, K. W., Gu, Z., Wang, Q., Sabuncu, M. R. & Kuceyeski, A. Release the Krakencoder: A unified brain connectome translation and fusion tool. Preprint at 10.1101/2024.04.12.589274 (2024).

16. Tozlu, C., Jamison, K., Gauthier, S. A. & Kuceyeski, A. Dynamic Functional Connectivity Better Predicts Disability Than Structural and Static Functional Connectivity in People With Multiple Sclerosis. Front. Neurosci. 15, 1683 (2021).

17. Boringa, J. B. et al. The brief repeatable battery of neuropsychological tests: normative values allow application in multiple sclerosis clinical practice. Mult. Scler. Houndmills Basingstoke Engl. 7, 263–267 (2001).

18. Goretti, B. et al. The brief international cognitive assessment for multiple sclerosis (BICAMS): normative values with gender, age and education corrections in the Italian population. BMC Neurol. 14, 171 (2014).

19. Skorve, E. et al. Brief international cognitive assessment for MS (BICAMS) and global brain volumes in early stages of MS – A longitudinal correlation study. Mult. Scler. Relat. Disord. 69, (2023).

20. Strober, L. et al. Symbol Digit Modalities Test: A valid clinical trial endpoint for measuring cognition in multiple sclerosis. Mult. Scler. Houndmills Basingstoke Engl. 25, 1781–1790 (2019).

21. Fuchs, T. A. et al. Repeated forms, testing intervals, and SDMT performance in a large multiple sclerosis dataset. Mult. Scler. Relat. Disord. 68, 104375 (2022).

22. Polman, C. et al. Diagnostic criteria for multiple sclerosis: 2010 revisions to the McDonald criteria. Ann. Neurol. 69, 292–302 (2011).

23. Tournier, J.-D. &, F. Calamante, and a. C. Improved probabilistic streamlines tractography by 2 nd order integration over fibre orientation distributions. Ismrm 88, 2010 (2010).

24. Tournier, J.-D., Calamante, F. & Connelly, A. MRtrix: Didusion tractography in crossing fiber regions. Int. J. Imaging Syst. Technol. 22, 53–66 (2012).

25. Tozlu, C., Card, S., Jamison, K., Gauthier, S. A. & Kuceyeski, A. Larger lesion volume in people with multiple sclerosis is associated with increased transition energies between brain states and decreased entropy of brain activity. Netw. Neurosci. 7, 539–556 (2023).

26. Fischl, B. & Dale, A. M. Measuring the thickness of the human cerebral cortex from magnetic resonance images. Proc. Natl. Acad. Sci. U. S. A. 97, 11050–5 (2000).

27. Tozlu, C. et al. Structural disconnectivity from paramagnetic rim lesions is related to disability in multiple sclerosis. Brain Behav. 11, e2353 (2021).

28. Tozlu, C. et al. The sequence of regional structural disconnectivity due to multiple sclerosis lesions. Brain Commun. 5, fcad332 (2023).

29. David, H. A. The beginnings of randomization tests. Am. Stat. (2008) doi:10.1198/000313008X269576.

30. Benjamini, Y. & Hochberg, Y. Controlling the False Discovery Rate: A Practical and Powerful Approach to Multiple Testing. J. R. Stat. Soc. Ser. B Methodol. 57, 289–300 (1995).

31. Yeo, B. T. T. et al. The organization of the human cerebral cortex estimated by intrinsic functional connectivity. J. Neurophysiol. 106, 1125–65 (2011).

32. Tian, Y. et al. Machine learning prediction of cognition from functional connectivity: Are feature weights reliable? bioRxiv 2021.05.27.446059 (2021) doi:10.1101/2021.05.27.446059.

33. Dhamala, E., Jamison, K. W., Jaywant, A. & Kuceyeski, A. Shared functional connections within and between cortical networks predict cognitive abilities in adult males and females. Hum. Brain Mapp. 43, 1087–1102 (2022).

34. Chen, J. et al. Shared and unique brain network features predict cognition, personality and mental health in childhood. 2020.06.24.168724 Preprint at 10.1101/2020.06.24.168724 (2020).

35. Seguin, C., Tian, Y. & Zalesky, A. Network communication models improve the behavioral and functional predictive utility of the human structural connectome. Netw. Neurosci. 4, 980–1006 (2020).

36. Finn, E. S. et al. Functional connectome fingerprinting: identifying individuals using patterns of brain connectivity. Nat. Neurosci. 18, 1664–1671 (2015).

37. Preziosa, P. et al. Structural connectivity in multiple sclerosis and simulation of disconnection (P4.367). Neurology 88, (2017).

38. Anderson, V. et al. A comprehensive assessment of cerebellar damage in multiple sclerosis using didusion tractography and volumetric analysis. Mult. Scler. J. 17, 1079– 1087 (2011).

39. Tozlu, C. et al. Structural disconnectivity from quantitative susceptibility mapping rim+ lesions is related to disability in people with multiple sclerosis. medRxiv 1–13 (2020).

40. Schmahmann, J. D. & Sherman, J. C. The cerebellar cognitive adective syndrome. Brain 121, 561–579 (1998).

41. Jacobi, H., Faber, J., Timmann, D. & Klockgether, T. Update cerebellum and cognition. J. Neurol. 268, 3921–3925 (2021).

42. Wenger, A., Calabrese, P. & Granziera, C. Unraveling the cerebellum’s role in multiple sclerosis. Curr. Opin. Behav. Sci. 56, 101357 (2024).

43. Schoonheim, M. M. et al. The cerebellum and its network: Disrupted static and dynamic functional connectivity patterns and cognitive impairment in multiple sclerosis. Mult. Scler. J. 135245852199927 (2021) doi:10.1177/1352458521999274.

44. Brouwer, E. J., Strik, M. & Schoonheim, M. M. The role of the cerebellum in multiple sclerosis: structural damage and disconnecting networks. Curr. Opin. Behav. Sci. 59, 101436 (2024).

45. Bonacchi, R. et al. The role of cerebellar damage in explaining disability and cognition in multiple sclerosis phenotypes: a multiparametric MRI study. J. Neurol. 269, 3841–3857 (2022).

46. Ziccardi, S. et al. Early regional cerebral grey matter damage predicts long-term cognitive impairment phenotypes in multiple sclerosis: a 20-year study. Brain Commun. 6, fcae355 (2024).

47. Weier, K. et al. Cerebellar Abnormalities Contribute to Disability Including Cognitive Impairment in Multiple Sclerosis. PLoS ONE 9, e86916 (2014).

48. Grothe, M., Domin, M., Hodeld, K., Nagels, G. & Lotze, M. Functional representation of the symbol digit modalities test in relapsing remitting multiple sclerosis. Mult. Scler. Relat. Disord. 43, 102159 (2020).

49. Bonavita, S. et al. Distributed changes in default-mode resting-state connectivity in multiple sclerosis. Mult. Scler. J. 17, 411–422 (2011).

50. Rocca, M. A. et al. Default-mode network dysfunction and cognitive impairment in progressive MS. Neurology 74, 1252–9 (2010).

51. Meijer, K. A. et al. Increased connectivity of hub networks and cognitive impairment in multiple sclerosis. Neurology 88, 2107–2114 (2017).

52. Thiebaut de Schotten, M., Foulon, C. & Nachev, P. Brain disconnections link structural connectivity with function and behaviour. Nat. Commun. 11, 5094 (2020).

53. Landmeyer, N. C. et al. Disease-modifying treatments and cognition in relapsing-remitting multiple sclerosis. Neurology 94, e2373–e2383 (2020).

