## Supplementary Document for "Predicting cognition using estimated structural and functional connectivity networks and artificial intelligence in multiple sclerosis"

### SUPPLEMENTARY INFORMATION

#### Network Modification Tool

The Network Modification 2.0 (NeMo 2.0) tool estimates the disruption in the structural connectivity due to a lesion or injury using a database of HC's structural connectivity. We first computed a database of whole-brain tractograms for 420 unrelated HC (206 female and 214 male, 28.7 ± 3.7 years) from the Human Connectome Project Young Adult (HCP-YA) dataset. The HCP-YA diffusion data has 1.25mm isotropic voxels, 3 shells (b=1000,2000,3000), and 90 directions per shell, and was collected with both R-L and L-R phase encoding. HCP data have been minimally preprocessed to correct for motion, EPI, and eddy-current distortion, and registered to subject T1 anatomy. We used MRtrix3 to estimate a voxel-wise multi-shell, multi-tissue constrained spherical deconvolution (CSD) model, followed by whole-brain deterministic (sd-stream) tractography with MRtrix3 with dynamic white-matter seeding. Streamlines for each HCP subject were warped into a common volumetric space (MNI152) to create the final reference set.

Supplementary Figure 1 shows that the models containing the pairwise eSC and eFC performed relatively worse than their regional counterparts

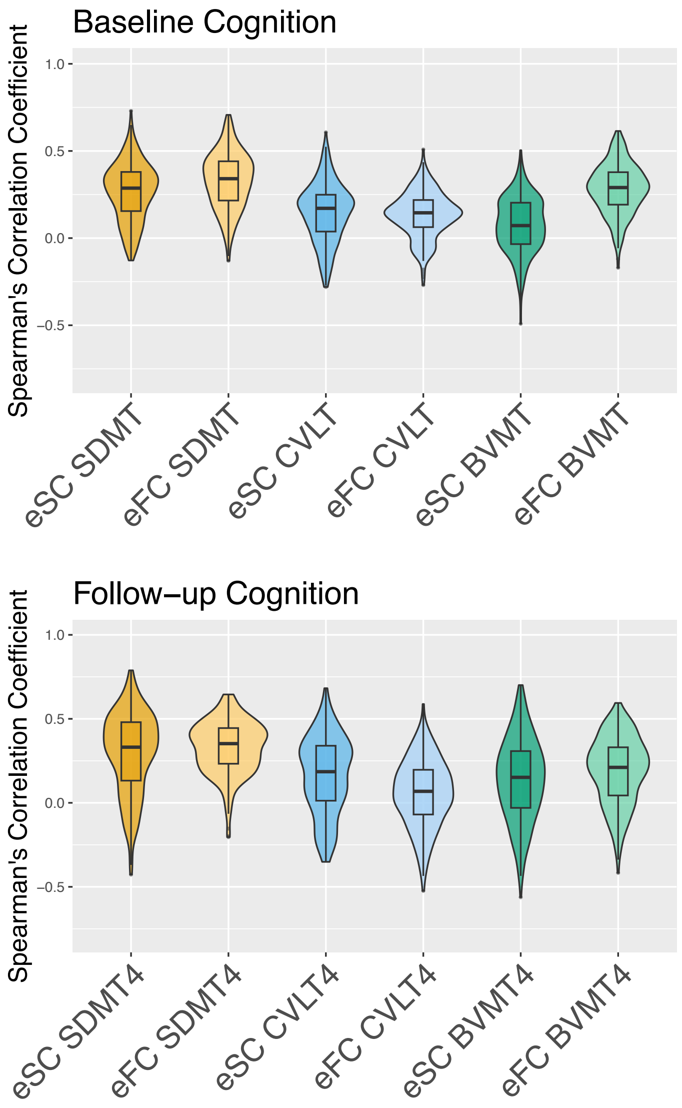


**Supplementary Figure 1**: The Spearman's correlation between the observed and predicted cognitive metrics obtained using models I and II where the regional eSC and eFC were replaced with their pairwise counterparts.

Supplementary Figure 2 shows the variable importance (beta estimates) in predicting baseline cognition. Among the models that included the connectomes, the highest accuracy was obtained with Model II which included demographics, clinical information, and clinic MRI neuroimaging features in addition to eFC in predicting baseline. Decreased eFC in the cerebellum was associated with higher baseline BVMT.

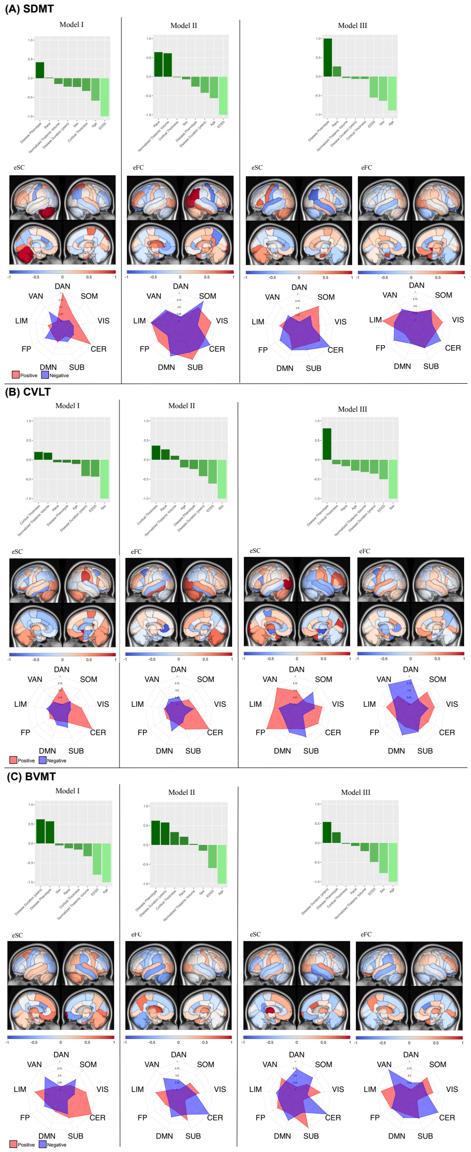

**Supplementary Figure 2**: Variable importance (beta estimates) of regional eSC, regional eFC, demographics, and clinical information as well as conventional MRI metrics calculated in 3 different models that were used to predict baseline SDMT, CVLT, and BVMT.

Supplementary Figure 3 shows the variable importance (beta estimates) in predicting follow-up cognition. Among the models that included the connectomes, the highest accuracy was obtained with Models I and II which included demographics and clinical information as well as conventional MRI metrics in addition to eSC and eFC, respectively in predicting follow-up SDMT. The interpretation of the variable importance was given in the main text.

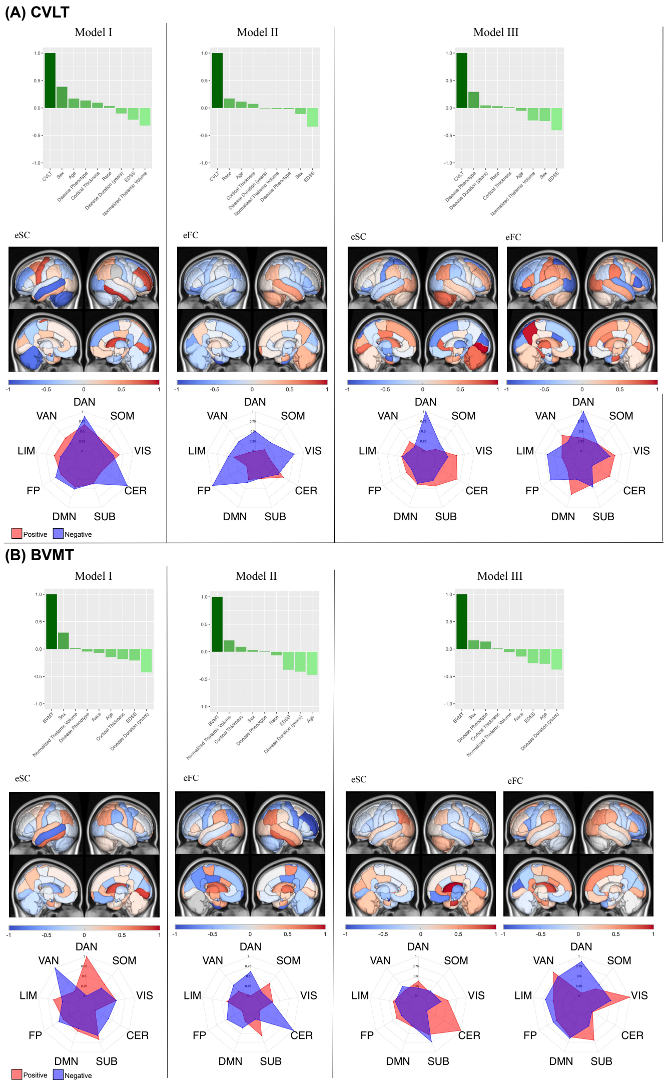


**Supplementary Figure 3:** Variable importance (beta estimates) of regional eSC, regional eFC, demographics, and clinical information as well as conventional MRI metrics calculated in 2 different models that were used to predict year 4 SDMT, CVLT, and BVMT.
